# The Advanced BRain Imaging on ageing and Memory (ABRIM) data collection: Study protocol and rationale

**DOI:** 10.1101/2023.11.16.567360

**Authors:** Michelle G. Jansen, Marcel P. Zwiers, Jose P. Marques, Kwok-Shing Chan, Jitse S. Amelink, Mareike Altgassen, Joukje M. Oosterman, David G. Norris

**Author notes:** Corresponding author (JMO). These authors contributed equally (Joukje M. Oosterman, David G. Norris).

## Abstract

To understand the neurocognitive mechanisms that underlie heterogeneity in cognitive ageing, recent scientific efforts have led to a growing public availability of imaging cohort data. The Advanced BRain Imaging on ageing and Memory (ABRIM) project aims to add to these existing datasets by taking an adult lifespan approach to provide a cross-sectional, normative database with a particular focus on connectivity, myelinization and iron content of the brain in concurrence with cognitive functioning, mechanisms of reserve, and sleep-wake rhythms. ABRIM freely shares MRI and behavioural data from 295 participants between 18-80 years, stratified by age decade and sex (median age 52, IQR 36-66, 53.20% females). The ABRIM MRI collection consists of both the raw and pre-processed structural and functional MRI data to facilitate data usage among both expert and non-expert users. The ABRIM behavioural collection includes measures of cognitive functioning (i.e., global cognition, processing speed, executive functions, and memory), proxy measures of cognitive reserve (e.g., educational attainment, verbal intelligence, and occupational complexity), and various self-reported questionnaires (e.g., on depressive symptoms, pain, and the use of memory strategies in daily life and during a memory task). In a sub-sample (*n* = 120), we recorded sleep-wake rhythms with an actigraphy device for a period of 7 consecutive days. Here, we provide an in-depth description of our study protocol, pre-processing pipelines, and data availability. ABRIM provides a cross-sectional database on healthy participants throughout the adult lifespan, including numerous parameters relevant to improve our understanding of cognitive ageing. Therefore, ABRIM enables researchers to model the advanced imaging parameters and cognitive topologies as a function of age, identify the normal range of values of such parameters, and to further investigate the diverse mechanisms of reserve and resilience.

## Introduction

Due to the worldwide ageing of the population, the proportion of adults who are suffering from or are at risk of cognitive impairment increases. This emphasizes the need to understand the building blocks of healthy cognitive ageing to reduce or even mitigate the decline in cognition that is associated with ageing [1, 2]. Normal ageing is accompanied by heterogeneous trajectories of cognitive decline within and across individuals, and predominantly affects processing speed, episodic memory, and reasoning [3, 4]. Where some individuals maintain a high level of cognitive functioning even throughout advanced age, others experience cognitive deficits that profoundly impact their daily lives [5, 6].

A growing body of research therefore seeks to explain individual variability in cognitive ageing by focusing on the underlying neural mechanisms [6, 7]. Brain ageing features that have been associated with lower cognitive functioning include a loss of brain volume and cortical thinning [8, 9], alterations in microstructural integrity, white matter organization, and cortical myelination [10–12], accumulation of iron content [13], and alterations in resting-state functional connectivity [14, 15]. However, patterns of brain ageing are highly variable across individuals, and marked differences exist in the extent of age-related brain changes as well as in the regional specificity of such alterations [16–18]. The concurrent investigation of brain ageing metrics therefore could provide complementary information on brain-cognition dynamics in ageing [18].

Another approach to study heterogeneity in cognitive ageing involves identifying the factors that contribute to risk or resilience [2, 6, 17, 19]. Factors such as hypertension, diabetes, obesity, physical inactivity, sleep disturbances, and depressive symptoms contribute to an increased risk of cognitive decline [2, 19]. In contrast, protective factors include, for example, educational attainment, occupational complexity, engagement in mentally stimulating activities, and social engagement [2, 17, 19]. It is assumed that risk factors of cognitive decline impact cognitive abilities through their associations with decreased brain health [20]. However, it remains to be elucidated whether protective factors promote cognition through neuroprotective effects (i.e., brain maintenance), stable neural advantages (i.e., brain reserve), and/or moderate the effects of neurodegeneration on cognition (i.e., cognitive reserve) [6, 17, 21, 22]. As brain health measures only partially explain heterogeneity in cognitive functioning, further insights on the precise working mechanisms of these protective factors are especially important [23].

Taken together, an extensive characterization of brain health parameters together with other factors of risk and resilience in cognitive ageing is warranted to increase our understanding of differing trajectories in ageing. Consequently, increasing efforts are made to facilitate the public availability of large neuroimaging datasets, directly via online repositories [24–27], or upon request and/or application [28–32].

The Advanced BRain Imaging on ageing and Memory (ABRIM) project aims to add to these existing datasets in several ways. First, only a few databases cover the adult lifespan [26, 29], whereas it is critical to map cognitive performance and brain health across all age groups. We therefore collected a cross-sectional, normative database of adults between 18-80 years old, stratified by age decade and sex. Second, quantitative imaging techniques are scarcely available in large population studies [24, 26, 31]. Quantitative imaging is particularly useful to investigate myelination and iron depositions in the brain [33–35], and has been associated with cognitive performance in normal aging [13, 36]. Our neuroimaging protocol therefore not only matches the sequences of the aforementioned datasets (i.e., conventional structural T1- and T2-weighted imaging, multi-shell diffusion weighted imaging, and resting-state functional MRI), but also facilitates quantitative imaging with Magnetization Prepared 2 RApid Gradient Echoes (MP2RAGE) and Multi Echo Gradient Echo Imaging (MEGRE) sequences. Where MP2RAGE allows to compute longitudinal relaxation rates (R_1_), MEGRE allows to derive apparent transverse relaxation rates (R_2_*) and quantitative susceptibility maps (QSM) [37, 38]. Lastly, in addition to cognitive test performance, we measured several variables associated with risk or resilience in cognitive ageing with self-reported questionnaires (e.g., on depressive symptoms and memory strategy use), a semi-structured interview (e.g., on educational attainment, occupational complexity, and leisure activities), and actigraphy (for sleep and physical activity estimates). This allows us to map the interactions of these variables with brain health across the lifespan more comprehensively. A complete overview of the variables that are included in ABRIM, in addition to the aforementioned neuroimaging datasets, is provided in S1 Table.

Below, we outline the study protocol of ABRIM, participant inclusion and exclusion procedures, behavioural and cognitive assessment, MRI sequences, and pre-processing pipelines. Furthermore, we describe our data sharing and management policies, as guided by the FAIR (Findable, Accessible, Interoperable, and Reusable) principles ([39]. ABRIM makes it possible to model imaging parameters and cognitive topologies throughout the adult lifespan, identify the normal range of values of such parameters, and further investigate the mechanisms that contribute to cognitive performance across the adult lifespan.

## Methods

### Participants

The study was performed at the Donders Institute for Brain, Cognition, and Behaviour, Radboud University Nijmegen the Netherlands. All data were collected between 2017–2022. The present study focused on a healthy, adult lifespan sample (18–80 years old).

We aimed to include approximately 25 male and 25 female participants per approximate age decade of adult life (18–30, 31–40, 41–50, 51–60, 61–70, 71–80). We recruited 301 participants from the general population, predominantly from the Nijmegen area. Several methods were employed to facilitate recruitment, including online and offline advertisements (e.g., on social media or in local supermarkets) and word of mouth. Participants were provided with a reimbursement of 10 euros per hour per bank transfer.

Potential participants were screened for inclusion via telephone or e-mail with a standardized questionnaire. Exclusion criteria consisted of the presence of any conditions with a profound impact on the brain and cognitive health, beyond normal aging, including current psychiatric disorders (e.g., major depressive disorder, bipolar disorder, schizophrenia), neurological conditions (e.g., dementia, history of stroke, epilepsy), substance use disorders (e.g., addiction to hard drugs), and history of other major health conditions that could impact cognition (e.g., history of brain tumour). Participants were additionally excluded if any MRI contraindications were present (e.g., ferromagnetic metal implants, claustrophobia, pregnancy).

Among the 301 participants tested, 6 participants were excluded from the dataset, either due to incidental MRI findings (*n =* 5) or because of the presence of a condition with a profound impact on the brain and cognitive health (*n = 1)*, resulting in 295 participants in the final study sample (53.20% females, median age 52 years, interquartile range [IQR] 36-66). An overview of the demographics of the ABRIM study per age decade is displayed in Fig 1. A more extensive description is provided in S2 Table. In a sub-sample (*n* = 119), we also acquired actigraphy data.

**Fig 1.**
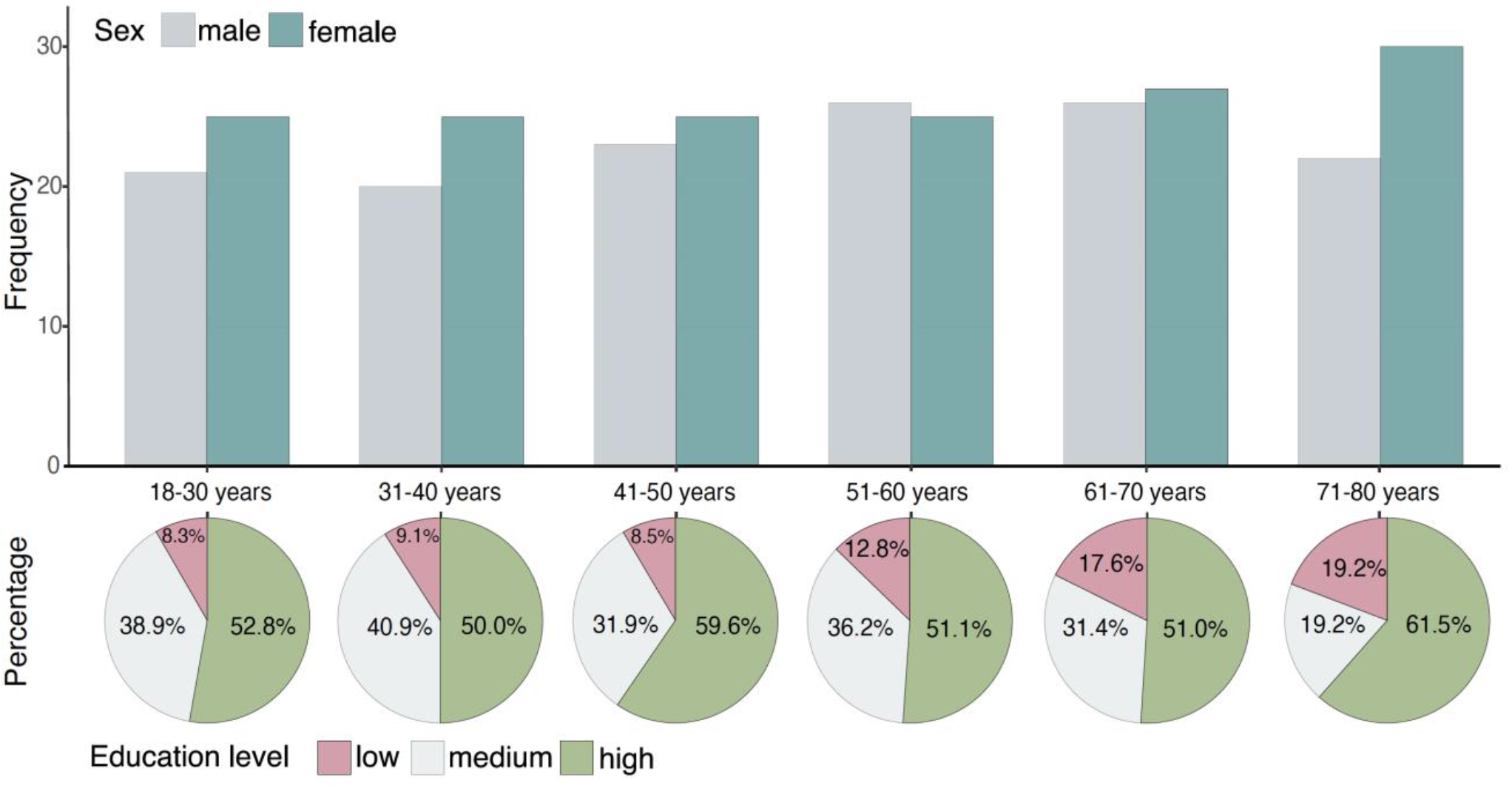
Demographics of the ABRIM study by age decade. The upper section displays the distribution of male and female participants for each age decade. The lower section consists of several pie charts, displaying the relative distribution of participants with low, medium, and high educational attainment within each age decade. Levels of education were classified using the International Classification of Education (ISCED-11): low education (ISCED 0-2), medium education (ISCED 3-5), and high education (ISCED 6-8). Data on educational attainment were not available for *n* = 18 participants.

Demographics of this sub-sample are displayed in S3 Table. Notably, we used the International Standard Classification of Education (ISCED-11 [40]) to classify educational attainment into three levels: low education (early childhood, primary, and lower secondary education, ISCED 0-2), medium education (upper secondary, post-secondary non-tertiary, and short cycle tertiary education, ISCED 3-5), and high education (bachelor, master, and doctoral or equivalent education, ISCED 6-8).

An additional n = 108 participants solely took part in the behavioural and cognitive assessment (sub-sample characteristics are available in S4 Table). These participants were included in the study prior to the start of the MRI acquisition, which was delayed due to a scanner upgrade. However, once the MRI acquisition was started (approximately 4-6 months after initial inclusion), these participants did not undergo the MRI protocol due to various reasons (e.g., no answer, lack of interest, unavailability, or MRI contraindications that were not previously mentioned). To ensure that our final study sample included approximately 300 participants, we extended our recruitment.

### Ethical approval

The entire study was conducted in compliance with the Helsinki declaration. The study fell under the blanket ethics approval “Image Human Cognition” (Commissie Mensgebonden Onderzoek Arnhem-Nijmegen, 2014/288), and was additionally approved by the Social Sciences Ethical Committee of the Radboud University (ECSW 2017-3001-46) and was conducted in compliance with all local procedures and applicable national legislation.

### Data management

We follow the requirements of the European General Data Protection Regulation (GDPR; https://gdpr.eu/) and the Dutch Act on Implementation of the General Data Protection Regulation.

To ensure anonymity of participants, privacy sensitive participant information (i.e., personal data) was separately stored in a password-protected database and only accessible to the main investigators. Participant data (i.e., scientific data) was anonymized and stored using study-specific numerical identification codes. A separate password-protected key file, that serves just to link participant identification codes to participant’s names, was kept by one researcher in strict confidence. We destroyed all privacy sensitive information and the key file one year after study completion, unless permission was granted by participants to be contacted in the future to be asked to participate in other studies. Signed informed consents and screening forms were locally archived in a closed locker at the Donders Institute, Nijmegen, the Netherlands, for at least 15 years after study completion. The original documents from the behavioural and cognitive examination (e.g., paper-and-pencil tests, self-reported questionnaires) were archived in a similar manner.

During data acquisition and analysis, raw actigraphy data was stored on a network directory from the Radboud University, Nijmegen, the Netherlands. With regards to MRI data, data were stored in a similarly secured way on network directory at the Donders Institute. Data storage is provided for by the main investigators of the study (MJ, JMO, DGN) and is accessible only for researchers involved in the processing of ABRIM.

With regards to data management of demographic variables and outcome measures of neuropsychological tests, self-reported questionnaires, and actigraphy, we used Castor (https://www.castoredc.com/). Castor provides a secured, cloud-based platform that supports researchers with adhering to Good Clinical Practice (GCP) guidelines (e.g., by saving all entered data and any changes that are being made). The study data is archived in Castor for 15 years for traceability purposes in accordance with GCP. All data stored in Castor is available for researchers involved in processing of the study after access has been granted by the main investigators.

For archiving purposes, raw MRI data, raw actigraphy data, and the Castor export were transferred to the Radboud Data Repository (RDR; https://data.ru.nl/). All research data is archived for at least 10 years after study completion.

Notably, the RDR consists of non-shared data collections for archiving purposes (i.e., data acquisition collection, DAC), as well as shared data collections of pre-processed or anonymized data (i.e., data sharing collection, DSC).

To facilitate data sharing and ensure pseudonymization, all anatomical scans were defaced prior to further processing, data was stored using non-identifiable codes, and data were minimized as much as possible (e.g., no birth data is shared, only age).

### Data sharing

ABRIM data is released through the RDR, with one DSC for the neuroimaging data (ABRIM MRI collection) and a second DSC for the cognitive and behavioural data (ABRIM behavioural collection).

Our data sharing procedures are in accordance with the agreements specified in the participants’ signed informed consent, and in consultation with the local privacy officer to ensure compliance to the relevant regulations, sharing as open as possible and as restricted as needed due to privacy legislation.

Neuroimaging data from ABRIM is estimated to be released in November 2023 (ABRIM MRI collection, https://doi.org/10.34973/7q0a-vj19, together with information on age and sex per subject (*n =* 295). We adhere to the guidelines outlined in the Brain Imaging Data Structure (BIDS; Gorgolewski et al., 2016), and COBIDAS MRI reporting framework [42]. A complete description of the measures in the ABRIM MRI collection is described below (“MRI”).

Cognitive and behavioural data will be released in November 2028 (ABRIM behavioural collection, https://doi.org/10.34973/7eq5-8y44). This data collection includes the measures obtained from neuropsychological tests, self-reported questionnaires, and actigraphy. Participants who solely participated in the cognitive and behavioural part prior to the start of the MRI phase are also included, resulting in a total of *n* = 404 participants. A complete description of the assessment protocol and measurements of the ABRIM behavioural collection is provided below (“Cognitive and behavioural data”).

Complete instructions on accessing the ABRIM MRI and behavioural DSCs will be provided on the RDR upon their release.

### Cognitive and behavioural data

#### Cognitive and behavioural examination

The cognitive and behavioural examinations were performed by trained researchers and consisted of a neuropsychological assessment, several self-report questionnaires, and a semi-structured interview to evaluate cognitive reserve.

The complete assessment took up to 2 hours and was performed in a quiet office-like environment without any distractions. Due to participant availability and logistics, not all assessments were performed on the same day as the MRI protocol (median days between assessments = 0, IQR 0-33.75). The order of the protocol was fixed to ensure that during the delay intervals for each memory test no new verbal stimuli to memorize were introduced. Below, we provide a detailed description of the different procedures and measures that were obtained. Notably, ongoing efforts to improve the cognitive and behavioural assessment introduced several novel components to the protocol throughout the course of the study, as indicated below.

#### Demographics, general health, and lifestyle variables

Standardized, self-report questionnaires were used to obtain information on demographics (age, sex, highest level of completed education), relevant medical conditions (e.g., hypertension, rheumatism), use of medication, and lifestyle habits (e.g., smoking behaviour, use of alcoholic beverages).

With regards to relevant medical conditions, participants were asked to report any current or previous psychiatric or neurological conditions, substance abuse, cerebrovascular accidents, cardiovascular disease, rheumatism, hypertension, hypercholesteremia, diabetes, sleep disorders, and chronic pain conditions. We evaluated the presence of these conditions not only to ensure compliance with the inclusion criteria, but also to inform on several, frequently occurring age-related conditions that did not warrant exclusion for the present study (e.g., rheumatism, hypertension, hypercholesteremia, diabetes, sleep disorders, chronic pain). In addition, participants were asked to rate their current and monthly pain using a numerical rating scale ranging between 0 (no pain) and 10 (worst pain imaginable).

Furthermore, participants were asked to provide a list of their current medication use. Subsequently, we classified them into the following categories: 1) psychoactive medication (e.g., antidepressants or antipsychotics); 2) tranquilizers or sleep medication (e.g., benzodiazepines); 3) anticoagulants (e.g., heparin); 4) antiplatelets (e.g., acetylsalicylic acid); 5) blood pressure medication (e.g., diuretics); 6) cholesterol medication (e.g., statins); and 7) diabetes medication (e.g., insulin).

We incorporated a customized, self-report questionnaire to the study after data were collected for ±100 participants to obtain information on smoking behaviour (current, ever smoking, or no smoking) and use of alcoholic beverages, in terms of average drinking frequency (never, monthly or less, 2-4 times a month, 2-3 times per week, 4 or more times per week), average number of alcoholic beverages when drinking (none, 1-2, 3-4, 5-6, 7-9, 10 or more beverages), and how often more than 6 alcoholic beverages are consumed (never, monthly or less, monthly, weekly, daily or almost daily).

#### Neuropsychological assessment

The neuropsychological examination consisted of a variety of tests measuring global cognitive functioning, processing speed, memory, and executive functions. In addition, we included a test to obtain an estimate of verbal intelligence.

##### Global cognitive functioning

The Montreal Cognitive Assessment (MoCA) was used as a measure of global cognitive functioning [43]. The MoCA consists of 11 items, tapping into several distinct cognitive domains. Visuospatial functions are measured using a cube- and clock-drawing task (4 points). Short-term memory involves learning 5 nouns and delayed recall after an interval of 5 minutes (5 points). Executive functions is measured using an alternation task (1 point), a phonemic fluency task (1 point), and a verbal abstraction task (2 points); attention, concentration and working memory by using a forward and backward digit-span task (2 points), a sustained attention task (1 point), and a serial subtraction task (3 points). Language was assessed using a three-item naming task with animals (3 points), the repetition of two sentences (2 points), and the beforementioned phonemic fluency task. Lastly, orientation to place and time is evaluated using different questions (e.g., “Tell me today’s date”; 6 points). As such, a total of 30 points can be obtained.

##### Verbal IQ

We used the Dutch Version of the National Adult Reading Test (DART) to obtain a measure of verbal IQ [44]. This test consists of a list of 50 written words that need to be read aloud by the participant that are scored by the experimenter based on pronunciation. To obtain a measure of verbal IQ, the raw scores are corrected for effects of age and sex, and subsequently transformed to indicate verbal IQ, based on Dutch norms. This test is often used as a proxy measure of CR, and may be a more sensitive measure of CR than education level [45, 46].

##### Memory functions

We evaluated memory functions using three different tests: The Story Recall subtest from the Rivermead Behavioural Memory Test – Third Edition (RBMT-3) [47], the Doors Test [48], and the Verbal Paired Associates (VPA) subtest of the Wechsler Memory Scale – Fourth Edition (WMS-IV-NL) [49]. Notably, the VPA was introduced after data was collected for ±100 participants already.

The RBMT-3 evaluates everyday memory functioning by using stimuli that correspond more to everyday life contexts compared to more traditional, laboratory tests of memory. During the Story Recall Subtest, the experimenter reads a 21-element story to the participant with the instructions to repeat as many items as possible afterwards (immediate recall). After an interval of about 15 minutes, the participant is again asked to recall as many elements as possible from this story (delayed recall). Completely correct elements are awarded one point, whereas partially correct elements are awarded half a point.

The Doors Test (part A and B) evaluates visual recognition memory [48]. For each part, participants are presented with 12 different target doors, each presented for 3 seconds. Immediately afterwards, the participant is asked to identify the target door on an array of 2 x 2 doors that includes 3 distractor doors. In part A, the distractors consist of different door types (e.g., a front door vs. a stable door, garage door, and café door). In contrast, distractors of visually similar door types are shown in part B (e.g., all front doors). Each correct response is rewarded with one point, and a total score of 24 can be obtained.

The VPA is a measure of associative memory [49]. The test consists of two parts. First, a list of 14 word-pairs is read to the participant that contains both semantically related (e.g., door-open) and semantically unrelated word pairs (plant-happy).

Subsequently, the experimenter provides the participant with the first word of a particular pair and asks the participant to recall the associated word. This procedure is then repeated three times, where the experimenter provides feedback on the participant’s responses (immediate recall). Second, after an interval of 20-30 minutes, the participant is asked to recall the paired words, again by providing the participant with the first word of a particular pair, but now without feedback from the experimenter (delayed recall). Subsequently, the delayed recall is followed by a yes/no recognition test of word pairs, and a free-recall test of words from the word pairs. During the free-recall test, the words can be recalled as single items as they do not necessarily have to be recalled within a particular pair. For each part of the VPA, we recorded the participant’s responses and the number of correct responses.

##### Executive functions and processing speed

We used three different tests to obtain measures for executive functions and processing speed: the Stroop Colour Word Test (SCWT [50]), the Trail Making Test (TMT [51]), and the digit span test of the WAIS-R (DST [52]).

The SCWT evaluates processing speed and inhibition [50]. Here, participants are asked to perform three tasks as fast as possible: 1) to read names of colours (Word Naming; W); 2) to name different colours (Colour Naming; C); 3) and to name the colour of the ink instead of reading the word itself while the colour-words are displayed in an incongruent colour (Colour-Word Naming; CW). For each task, we recorded the completion time and the number of errors made.

The TMT consists of part A and part B, and allows to measure attention and processing speed [53]. In part A, participants connect a set of 25 consecutive numbers. In part B, a set of 25 circles is connected by alternating between numbers and letters. During both parts, participants are instructed to work as fast and accurately as possible. For each part, the time to complete the task and the number of errors is recorded.

We used the DST to measure working memory. The experimenter reads a list of numbers and asks the participant to recall this list immediately afterwards in forward order. Subsequently, a second list is presented, where the participant is asked to recall the numbers in backward order. For both lists, the total number of digits increases after each sequence of two trials, until the participant fails a complete sequence or reaches the end of the test list that each consists of 6 sequences. The total number of correct reproductions on the forward and the backward versions are recorded.

#### Cognitive reserve index questionnaire

We used the Cognitive Reserve Index questionnaire (CRIq) to obtain an indication of CR. The CRIq is a semi-structured interview focusing on several proxy measures of CR, namely: education, working activity, and leisure activities. These domains have been suggested to predominantly contribute to the lifetime experiences that contribute to CR [5, 54]. The CRIq hence does not capture CR directly, but instead provides an overall indication of CR that has been accumulated throughout the lifespan as well as sub-scores for each separate domain [54]. Previous literature demonstrated that CRIq scores indeed could explain the discrepancy between cognitive performance and brain pathology in various study populations (i.e., from healthy to pathological ageing), indicating that this is a valid indicator of CR [5, 55].

#### Self-report questionnaires

Self-report questionnaires included the Beck Depression Inventory, Brief Pain Inventory, Self-Report Psychopathy-short form, and the Metamemory in Adulthood Questionnaire-short form [56–61]. Notably, as mentioned earlier, ongoing efforts to improve the cognitive and behavioural assessment introduced several novel components to the protocol throughout the course of the study. After the inclusion of ±100 participants, the Everyday Memory Questionnaire (revised), Cognitive Failure Questionnaire, and a customized strategy use questionnaire were added to our protocol [62–64].

##### Beck Depression Inventory (BDI)

The BDI was implemented to identify potentially high levels of depressive symptoms and evaluates depressive symptoms on a 4-point scale across 21 items, where higher scores indicate increased presence of depressive symptomatology [56]. A score between 0-9 indicates that an individual is experiencing no or minimal depression, 10-18 indicates mild depression, 19-29 indicates moderate depression, and 30-63 indicates severe depression.

##### Brief Pain Inventory (BPI-SF)

The BPI-SF measures presence and location of pain, pain severity, pain interference on daily functions, pain treatment and (the percentage of) pain relief following treatment. Pain severity (4 items) and pain interference (7 items) are rated on an 11-point rating scale ranging from 0 to 10 [57, 58].

##### The Self-Report Psychopathy – Short Form (SRP-SF)

The SRP-SF evaluates psychopathy-related traits [59]. This questionnaire consists of 28 items, rated on a 5-point scale, and comprises four subscales: Interpersonal Manipulation, Callous Affect, Erratic Lifestyle, and Criminal Tendencies.

##### Metamemory in Adulthood questionnaire – Short Form (MIA-SF)

The MIA-SF evaluates subjective memory functions and knowledge of memory processes [60]. In the present study, we use the abridged Dutch version of this questionnaire [61]. The questionnaire consists of 74 items, rated on a 5-point scale, and allows to calculate separate scores for the following domains: Task, Capacity, Change, Anxiety, Achievement, Locus and Strategy. The Strategy domain contains 16 items and is additionally divided into External Strategies (e.g., memory aids, 8 items) and Internal Strategies (e.g., mental imagery, 8 items).

##### Everyday Memory Questionnaire - Revised (EMQ)

The EMQ measures subjective memory failure in everyday life [64]. This questionnaire consists of 13 items, rated on a 5-point scale. Each item focuses on the frequency of memory failures in daily life, such as forgetting when a certain event happened (e.g., whether this occurred yesterday or last week), or forgetting to tell someone something important (e.g., passing a message from someone else).

##### Cognitive Failure Questionnaire (CFQ)

The CFQ evaluates subjective cognitive functioning and consists of 25 items, rated on a 5-point scale [62, 63]. Each item focuses on the frequency of daily cognitive mistakes, such as missing appointments or experiencing difficulties in making decisions. Four additional questions inform on potential increases in the occurrence of these mistakes and the extent to which an individual finds these experiences troublesome, annoying, or worrisome; however, these items are not used in scoring this questionnaire.

##### Strategy use

We incorporated a separate, custom-made questionnaire to obtain information on strategy use during the VPA. Here, after completing the VPA, participants are asked to describe whether they used strategies to recall the word pairs, and the type of strategies they used (e.g., concentrating, repetition, visualization, association; see S1 Appendix for an English translation of this questionnaire).

### Actigraphy

We used the Actiwatch 2 (Philips Respironics, Eindhoven, the Netherlands), a wristband-type actigraphy device with an internal accelerometer and light sensor to infer sleep-wake rhythms. As we did not have enough devices to acquire actigraphy data among all participants, for only a sub-sample of the participants we were able to record actigraphy data (*n* = 134). The decision to give a participant an actiwatch was based on the availability of the devices, while also aiming to achieve an equal age distribution of this sub-sample. Actigraphy was acquired for 7 consecutive days.

Participants were asked to report any sleep disorders, as these could affect the data collection (e.g., insomnia, sleep apnea, restless leg syndrome). Epoch length was set at 15 seconds. All data was processed with Actiware software (v6.0.9).

All actograms were visually checked to ensure quality of the data. Data were excluded when the algorithm failed to register any sleep/wake rhythms or was unable to delineate these (e.g., because the device was not worn at all or too irregularly). A minimum of 5 days of usable data was required for inclusion of the data in our database. This resulted in the exclusion of *n* = 14 participants (*n = 6* failed to register any activity, *n =* 3 unable to properly distinguish sleep/wake rhythm, *n* = 5 with less than 5 days of usable data). Therefore, a total of *n* = 120 participants were included in our database. Data of an additional *n* = 11 participants were acquired amongst those who only participated in the cognitive and behavioural measurements (*n* = 1 failed to register any activity and was excluded, resulting in *n* = 10 participants).

With regards to sleep, we recorded the following outcomes measures: total sleep time (summation of sleep epochs within the sleep phases), wake after sleep onset (WASO; summation of wake epochs between begin and end of a sleep phase), sleep latency, sleep efficiency, and number of awakenings. Although this device has been predominantly shown to measure sleep in an objective and reliable manner [65], recent studies demonstrated that valid measures of physical activity can also be calculated, for example, by using the activity counts per minute or cycle recorded by the device [66].

### MRI

#### Data acquisition

All scans were acquired on a 3T Siemens Magnetom Prisma System (Siemens, Erlangen, Germany) using the standard 32-channel receive coil. We used an auto-align localizer sequence to automatically align all imaging sequences, additionally each alignment was visually checked and manually adjusted when necessary. The duration of the complete scanning session was approximately 55 minutes. Participants were held with head cushions and were additionally fixed using a small piece of tape to reduce head movement [67].

The following MRI scans were acquired: 1) T1-weighted 3D Magnetization Prepared - RApid Gradient Echo (MPRAGE); 2) T1-weighted MP2RAGE; 3) Fast turboFLASH B1 mapping; 4) T2-weighted turbo spin echo (TSE) sequence; 5) multi-shell High Angular Resolution Diffusion Imaging (HARDI); 6) Multi-echo gradient echo (MEGRE); and 7) resting-state BOLD fMRI (GE-EPI). A complete overview of the sequence parameters can be found in Table 1 and a more detailed account in the BIDS metadata files in the *bids* subject folders (part of the ABRIM MRI collection).

**Table 1.** Overview of neuroimaging sequences and parameters

| Scan type | Sequence | TR<br>(ms) | TE<br>(ms) | Flip<br>angle<br>(°) | Voxel size<br>(mm) | FOV<br>(RL/AP/<br>FH) | Other |
| --- | --- | --- | --- | --- | --- | --- | --- |
| T1-weighted | MPRAGE | 2200 | 2.64 | 11 | 0.8x0.8x0.8 | 180x320x<br>256 | GRAPPA: 3;<br>TI = 1100 |
|  | MP2RAGE | 6000 | 2.34 | 6/6 | 1.0x1.0x1.0 | 263x350x<br>350 | GRAPPA: 3;<br>TI =<br>700/2400 |
| B1 map | turboFLASH | 10000 | 2.23 | 8 | 3.3x3.3x2.5 | 350x263x<br>350 |  |
| T2-weighted | TSE | 3200 | 569 | n/a | 0.8x0.8x0.8 | 180x320x<br>256 | GRAPPA: 3;<br>variable flip<br>angle |
| Diffusion-weighted | PGSE-EPI | 2940 | 74.8<br>0 | 90 | 1.8x1.8x1.8 | 216x216x<br>146 | GRAPPA: 2;<br>SMS factor =<br>3<br>(interleaved);<br>11 b=0 s/mm<sup>2</sup><br>volumes; 86<br>b=1250<br>s/mm<sup>2</sup>; 85<br>b=2500<br>s/mm<sup>2</sup> |
| MEGRE | Bipolar<br>multi-echo<br>GRE | 44 | TE<sub>1</sub>/<br>DTE<br>/<br>TE<sub>9</sub><br>=<br>6.14/ | 20 | 0.8x0.8x0.8 | 186x230x<br>141 | GRAPPA: 3 |
|  |  |  | 4/38.<br>14 |  |  |  |  |
| Resting-state fMRI | Multi-band<br>GRE-EPI | 1000 | 34 | 60 | 2.0x2.0x2.0 | 210x210x<br>132 |  |
|  | Inverted<br>Distortion<br>GRE-EPI | 1000 | 34 | 60 | 2.0x2.0x2.0 | 210x210x<br>132 |  |

TR, repetition time; TE, echo time; TI, inversion time; FOV, field of view; MPRAGE,
Magnetization Prepared Rapid Acquisition Gradient Echo; GRAPPA, GeneRalized Autocalibrating Partial Parallel Acquisition; MP2RAGE, Magnetization Prepared 2 Rapid Acquisition Gradient Echoes. TSE, Turbo Spin Echo. PGSE, Pulsed-gradient spin-echo. EPI, T2\*-weighted gradient echo planar imaging. SMS, simultaneous multi-slice. GRE, Gradient Echo Imaging.

Furthermore, among *n* = 76 participants, task-based fMRI was collected during a prospective memory paradigm, together with the prospective and retrospective memory questionnaire [68–70]. However, as the prolonged duration of the total MRI scanning protocol (±2.5 hours) led to significant recruitment difficulties, the acquisition of the task was discontinued to preserve the feasibility of the study.

Several efforts were made to facilitate the shareability, reproducibility, and reusability of ABRIM MRI data [71]. First, the ABRIM MRI collection adheres to the BIDS standard (Gorgolewski et al., 2016). Second, we used standard containerized neuroimaging pipelines specifically developed for BIDS data (i.e., BIDS apps, Gorgolewski et al., 2016) to process the raw MRI images and to provide visual quality control (QC) reports. Below, we explain our MRI sequences and (pre-)processing pipelines in more detail. A schematic illustration of the various methodological steps that were applied to each imaging modality, and corresponding BIDS folders, is displayed in Fig 2. An overview of the BIDS folder structure itself is provided in S1 Fig.

**Fig 2.**
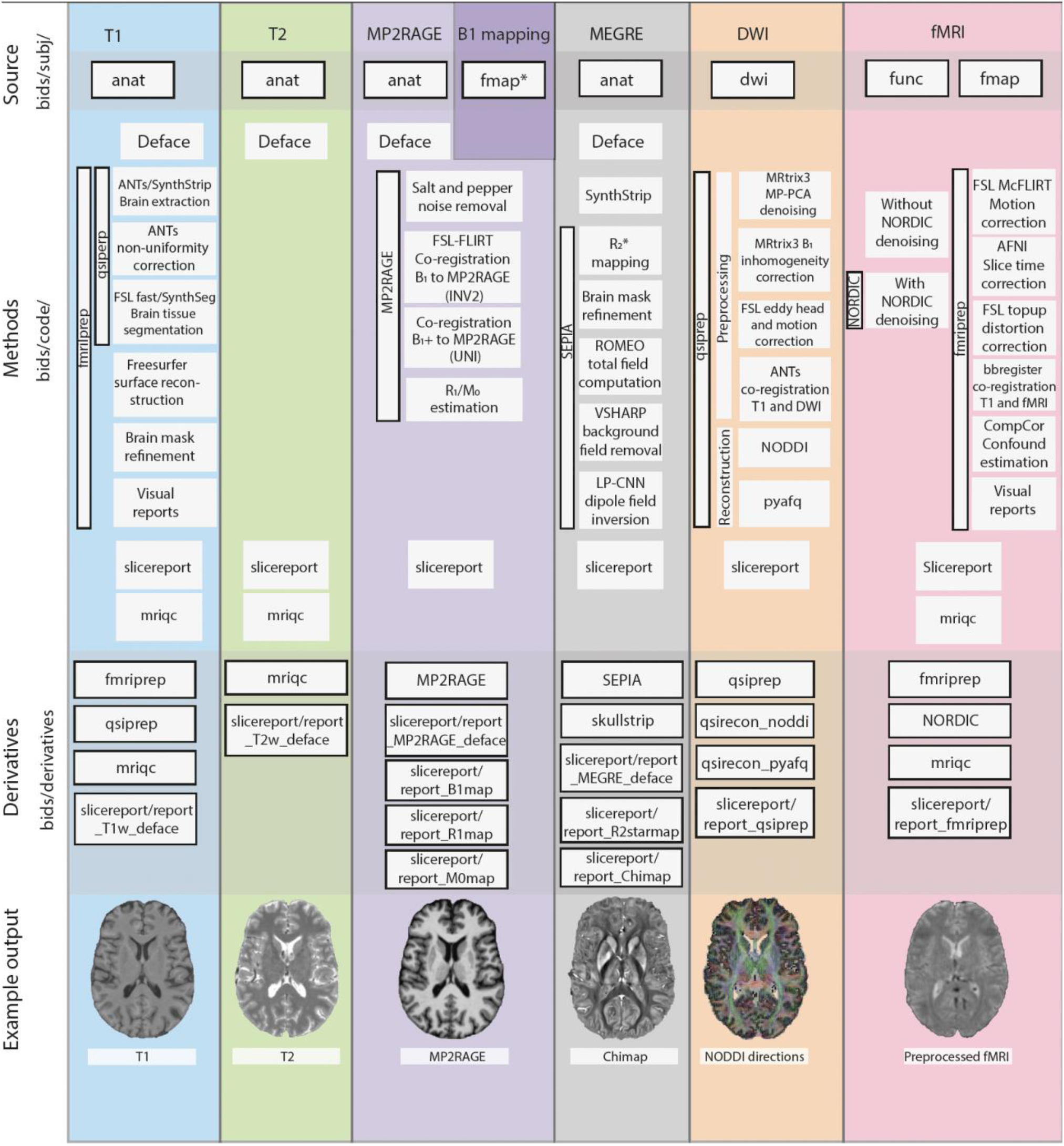
Methodological steps applied to each imaging modality in ABRIM. Schematic illustration of the methodological steps that were applied for each imaging modality, and its corresponding Brain Imaging Data Structure (BIDS) folders. Each column in the figure represents a different imaging modality, and is distinguished by a unique colour (e.g., T1 images in blue). Each row represents a specific methodological step (e.g., fMRIprep) with its corresponding sub-steps (e.g., brain extraction). BIDS folders are denoted with black frames. The process starts with the source data (e.g., bids/subj/anat/ folder) and progresses to the derivatives (e.g., bids/derivatives/fmriprep/ folder). For each imaging modality, an example output image is displayed at the bottom. All code utilized can be found in the bids/code folder. Note that a certain imaging modality may be processed by multiple methods (e.g., T1 images with both fMRIprep and QSIprep). *fmap is available under bids/subj/derivatives/SIEMENS/

#### BIDS conversion

To transform raw DICOM files to BIDS, we used version 3.7.4 of the open-source BIDScoin application (https://github.com/Donders-Institute/bidscoin) [72]. The resulting data complied to version 1.8 of the BIDS standard, which was verified by application of the bids-validator version 1.11 (https://github.com/bids-standard/bids-validator) After BIDS conversion was completed for all participants, we proceeded with de-identification of all anatomical MRI data by applying the deface BIDS app of BIDScoin. The deface app is a wrapper of *pydeface* (https://github.com/poldracklab/pydeface) and ensures that the output corresponds to the BIDS standard. All output was visually inspected by one author (MGJ) to ensure successful de-identification, while a second author was consulted (MPZ) in case of any uncertainties. Manual masks were created in case of any remaining unique personal features that could led to potential identification (e.g., nasal features), and masked out from the anatomical images using *fslmaths* from FMRIB Software Library (FSL; https://fsl.fmrib.ox.ac.uk/).

All pre-processed MR images and QC reports (as described in more detail in the sections below), are available in the *bids/derivatives* folder, and the scripts to generate them in the *bids/code* folder (both part of the ABRIM MRI collection).

#### T1- and T2-weighted imaging derivatives

We incorporated the T1-weighted MPRAGE and T2-weighted TSE sequences to allow for the characterization of conventional markers of brain ageing, such as the volume and thickness of various brain regions [73, 74]. Furthermore, these sequences can be combined to obtain more detailed measurements of brain morphology and maps of the structural organization of the cerebral cortex (e.g., relative myelin content) [75]. As explained below, the T1 images are minimally processed together with the resting-state fMRI sequence. For both T1 and T2 images, we provide visual QC reports.

We used the general purpose slicereport-tool of BIDScoin to generate visual QC reports unless otherwise specified. Slicereport facilitates the generation of a web page, displaying rows of image slices for each subject, and optionally provides individual sub-pages with more detailed information. In addition, it provides the flexibility to customize the displayed information. Therefore, slicereport enables efficient and thorough visual inspections of MRI data. For an example of its application in ABRIM, see Fig 3.

**Fig 3.**
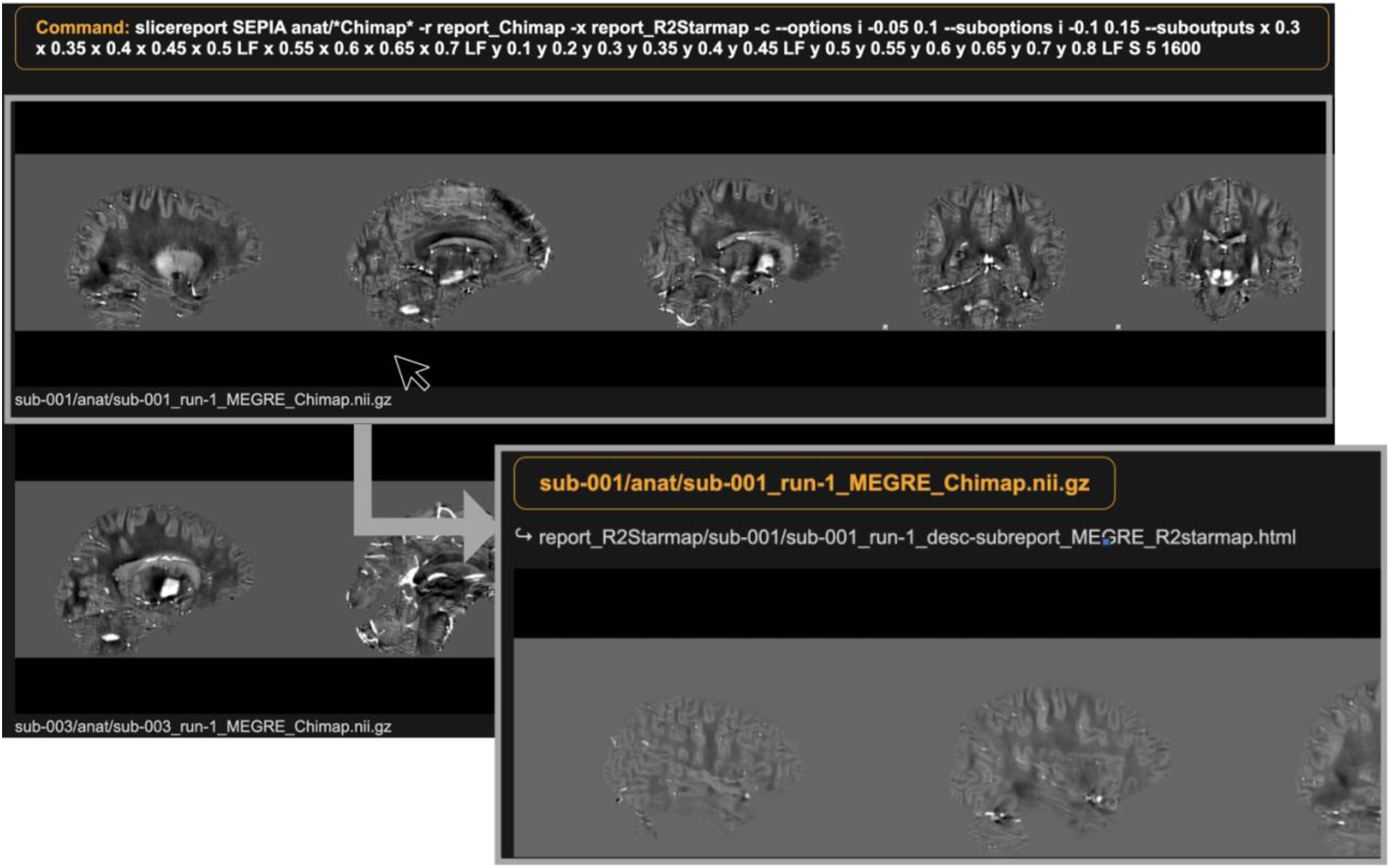
Example of slicereport output for ABRIM. An example of the slicereport-tool to create visual quality control reports for Quantitative Susceptibility Mapping (QSM) output in ABRIM. The top left image displays the generated web page, featuring rows of image slices per subject. In this case, the researchers opted for several sagittal, coronal, and axial slices to enable visual inspection of physiological noise (e.g., motion, ringing) and streaking artifacts. Upon clicking on the image slices of a specific subject, it leads to a sub-page (right bottom image). In this case, the sub-page displays corresponding image slices from the R_2_* maps to facilitate easier delineation of motion artifacts.

Asides from this, we applied the MRI Quality Control tool (MRIQC) to provide additional QC reports and image quality metrics for the T1, T2, and fMRI images [76]. Corresponding group reports are available in S2 Fig, S3 Fig, and S4 Fig.

#### MP2RAGE and Fast turboFLASH B1 mapping

The MP2RAGE sequence was developed to obtain T1-weighted images that are less dependent of transmit and receive field inhomogeneities (B_1_ ^+^ and B_1_ ^-^) and improve the contrast between white matter and grey matter tissue [37] by removing residual dependency on water density and transverse relaxation (M_0_ and T_2_*). We acquired MP2RAGE in combination with a turbo flash B_1_ map to obtain quantitative longitudinal relaxation rate (R_1_) maps [33] as well as proton density maps. Previous studies revealed that R1 maps are useful to delineate cortical myelination in group studies [35], and to study white matter tissue maturation over the lifespan [77]. Proton density maps can be used to compute the Macromolecular Tissue Volume fraction [78].

To process the MP2RAGE images, we used BIDS compatible Matlab scripts that are hosted on Github (https://github.com/Donders-Institute/MP2RAGE-related-scripts). For the purpose of anatomical referencing, the typical salt and pepper noise of the MP2RAGE was reduced using the regularization method as described in [79].

Quantitative R1 maps were obtained using bids_T1B1correct.m from the above repository: (i) The magnitude image from the turbo flash B1 map was co-registered to the MP2RAGE image with proton density contrast (INV2) using SPM12 [80]; (ii) The resulting transform matrix was then used to co-register a spatially smoothed version of the B1+ map to the MP2RAGE image; (iii) To correct for B1 transmit inhomogeneities in R1 estimations (and derive M0 maps over a large range of R1 values), instead of the traditional MP2RAGE lookup table, a fingerprinting-like approach was used to estimate R1 [81]. Here, the maximum of the inner product of the signal at the two inversion times is used by a dictionary to determine the voxel R_1_ and M_0_. MP2RAGE signal dictionaries were computed at steps 0.005 nominal B_1_ field. Finally, visual QC reports were generated for the R_1_ and M_0_ maps.

#### Gradient echo imaging

We used MEGRE for apparent transverse relaxation rate (R_2_*) mapping and quantitative susceptibility mapping (QSM) to obtain quantitative measurements that relate to local concentrations of iron and myelin in the brain. Both techniques are sensitive to the change of tissue magnetic susceptibility *(𝜒)*. Conventionally, the magnitude of the MEGRE data is used to derive R_2_* maps and the phase of the MEGRE data is used for QSM. Previous studies demonstrated that increases of iron and myelin concentration enhance the R2* values [38, 82, 83]. In QSM, however, these two constituents produce opposite image contrasts since iron is paramagnetic (+𝜒) and myelin is diamagnetic (-𝜒) with respect to water [38, 82, 84]. As ageing is characterized by changes in both myelination and iron depositions throughout the brain [34, 85], processing both the QSM and R_2_*\** maps provides a more complete picture of these alterations at no extra cost as both maps can be derived from the same data.

We used custom BIDS wrapper code around the SEPIA toolbox (v1.2.2.3) for generation of the R_2_* and QSM maps [86]. Prior to applying the SEPIA toolbox, we acquired brain masks using SynthStrip, a novel learning-based brain extraction tool with highly accurate performance across different imaging modalities and populations [87]. More specifically, we implemented SynthStrip using the skullstrip BIDS app from BIDScoin.

R_2_* maps were derived by extracting the magnitude of the MEGRE data based on a closed-form solution [86, 88]. Additionally, we generated the corresponding visual QC reports.

We acquired QSM maps through multiple steps in SEPIA [86]. First, the MEGRE brain masks were initially refined by masking out high R2* voxels on the mask edge associated with non-tissue of interest (e.g., air and vein) that can create strong susceptibility artefacts. Subsequently, total field computation was performed with ROMEO [89], background field removal with VSHARP [90], and dipole field inversion using LP-CNN [91]. As QSM only provides relative values, we used the mean susceptibility of the whole brain as a reference to facilitate data comparison between subjects. Visual QC reports were generated for the QSM maps.

In addition to the QC reports, we provide subjective quality assessment ratings for physiological noise (e.g., motion, ringing) and streaking artifacts in QSM. Both aspects can have a profound impact on the contrast of the resulting images and validity of results [92]. The presence of both quality aspects was visually rated on a scale from 0 (optimal quality) to 5 (multiple / gross artefacts). Subsequently, these scores are summed to indicate the total quality. The quality assessment file also includes a dichotomous rating to indicate if any parts of the cerebellum or cerebrum were missing (0 = complete; 1 = missing).

#### Diffusion-weighted imaging

Diffusion-weighted imaging allows to investigate the microstructural properties of the brain in a non-invasive manner. We used multi-shell HARDI with 182 diffusion directions (HARDI, 11 x b = 0, 86 x b = 1250, 85 x b = 2500 s/mm^2^) to derive several measures, such as fractional anisotropy (FA), mean diffusivity (MD) [93] or orientation dispersion (OD) [94], reflecting different microstructural properties.

HARDI also allows for fibre tractography, where diffusion-derived measures are modelled along the trajectories of white matter pathways [95]. Age-related differences and neurodegenerative diseases have been frequently related to these metrics, where loss of white matter integrity is typically characterized by decreased FA and increase MD values [96, 97].

It is well-known that diffusion imaging data are sensitive to corruption from various sources of physiological and scanner noise, and that the images are typically geometrically distorted due to local (susceptibility-induced) and global (eddy-current-induced) magnetic field distortions [98]. We used the state-of-the-art BIDS compatible QSIPrep pre-processing pipeline (v0.18.0) to correct for artifacts and to offer high-quality data.

In short, QSIprep processes both structural MRI and diffusion MRI, and automatically incorporates pre-configured workflows based on the data provided to estimate various popular HARDI models, QC metrics, and visual QC reports [99]. For ABRIM structural MRI, QSIprep uses Advanced Normalization Tools (ANTs, https://www.nitrc.org/projects/ants) to correct for intensity non-uniformity [100] and for non-linear registration of T1 images to the MNI152 template [101]. Brain extraction is performed using SynthStrip [87], and tissue segmentation using SynthSeg [102]. For ABRIM diffusion data, QSIprep applies MP-PCA denoising as implemented in MRtrix3’s dwidenoise [103], B1 field inhomogeneity correction using dwibiascorrect from MRtrix3 (https://www.mrtrix.org/) with the N4 algorithm [100], head motion and eddy current correction using FSL’s eddy [104], and co-registration to the T1-weighted image using antsRegistration (ANTs) [101]. The complete QSIprep pre-processing configuration is described within the ABRIM MRI collection (*/bids/derivatives/qsiprep/logs*).

We subsequently used QSIprep to apply two preconfigured reconstruction workflows. The first workflow (*mrtrix_multishell_msmt_pyafq_tractometry*) distinguishes major white matter pathways and estimates its corresponding tissue properties. More specifically, besides providing FA and MD estimates, this workflow applies automatic fiber-tract quantification (AFQ [105]) [106] and uses IFOD2 from MRtrix3 to generate many more microstructural measures and tractography data [107]. The second workflow (amico_noddi) was utilized to estimate the neurite orientation dispersion and density imaging (NODDI) model [94] with the AMICO implementation [108]. The resulting outputs include the intra-cellular volume fraction (ICVF), isotropic volume fraction (ISOVF), and orientation dispersion (OD).

#### Resting-state fMRI

Resting-state fMRI was incorporated because of its relation to task-activation, brain network organization and cognition [109, 110]. With advancing age, brain networks are characterized by less distinct functional networks and decreased local efficiency, predominantly among the brain networks responsible for higher order cognitive processes [14].

We applied the BIDS compatible fMRIprep pre-processing pipeline (v23.1.2), which facilitates the pre-processing of structural and fMRI data by combining different software packages [111]. Briefly, T1 images are corrected for intensity non-uniformity and brain extracted using ANTs [100]. Brain surfaces are reconstructed using Freesurfer (v7.3.2. https://surfer.nmr.mgh.harvard.edu/) and used to refine the initially estimated brain mask. Non-linear registration of T1 images to the ICBM152 template is performed with ANTs [101], and brain tissue segmentation with FSL FAST [112]. With regards to fMRI, susceptibility-induced distortion corrections are performed using FSL topup [113], slice time corrections with Analysis of Functional NeuroImages (AFNI, https://afni.nimh.nih.gov/), and motion corrections with FSL McFLIRT [114]. Co-registration between the T1 images and fMRI data is facilitated using bbregister from Freesurfer [80]. Confounds are estimated for nuisance regression using CompCor [115]. In addition, visual QC reports are saved that allow for a complete evaluation of the fMRIprep procedures. The complete fMRIprep pre-processing configuration is described within the ABRIM MRI collection (bids/derivatives/fmriprep/logs).

In addition, we applied Noise Reduction with Distribution Corrected (NORDIC) denoising on the fMRI data, using both the magnitude and phase time series (https://github.com/SteenMoeller/NORDIC_Raw), before running the fMRIprep pre-processing pipeline on the denoised data. Briefly, NORDIC allows to further improve the spatial temporal resolution of fMRI data by reducing thermal noise (i.e., white noise due to thermal fluctuations of the subject and/or receive coil). In contrast to more traditional denoising algorithms (e.g., ICA-AROMA [116]), NORDIC does not affect structured, non-white noise (e.g., due to respiration or the cardiac cycle), and therefore complements other algorithms [117, 118].

## Discussion

To increase our understanding of why we all age differently, an extensive characterization of brain health in concurrence with other factors of risk and resilience in cognitive ageing is warranted [2, 6, 17, 19]. Conversely, this has led to the growing availability of large datasets to study the dynamics of cognitive ageing [24–32].

With ABRIM, we aimed to add to existing study cohorts by 1) collecting a cross-sectional, normative database of adults between 18-80 years old, stratified by age and sex; 2) incorporating both conventional imaging sequences as well as quantitative imaging methods; 3) concurrently measuring cognitive functions with mechanisms of reserve (e.g., cognitive reserve), and actigraphy (e.g., to establish sleep-wake rhythms). The present study provides a publicly shared cross-sectional data repository on healthy adults throughout the lifespan, including numerous parameters relevant to improve our understanding of cognitive ageing. More specifically, the ABRIM MRI collection includes BIDS-compliant raw and pre-processed MRI measures and derivatives of T1- and T2-weighted imaging, quantitative imaging (MP2RAGE and gradient echo imaging), multi-shell diffusion-weighted imaging, and resting-state fMRI. In addition, all code is provided that generated the derivatives. The ABRIM behavioural data collection complements these MRI measures with the outcomes of a neuropsychological examination (e.g., on executive functioning, processing speed, memory, and global cognition), self-reported questionnaires (e.g., on lifestyle factors, depression, pain, psychopathy, memory strategy use), and actigraphy measures (e.g., sleep-wake parameters).

Nevertheless, there are several limitations that need to be addressed. First, the generalizability of our sample is limited as participants are relatively healthier and more highly educated as compared to the general population. Furthermore, we only covered the adult lifespan between 18-80 years old to minimize potential confounding factors associated with extreme age ranges and for feasibility purposes. Consequently, our sample is not fully reflective of the ageing population. In addition, as ABRIM is a cross-sectional database, this merely allows us to examine cognitive performance within a specific age range at a single point in time, while mapping of individual trajectories of cognitive decline is impossible. Lastly, our sample size remains relatively small as compared to other publicly available datasets. Therefore, we emphasize the importance of replicating findings across multiple cohort studies, and hope that the ABRIM dataset will further encourage open and reproducible research in the field of cognitive ageing [71].

## Conclusion

Altogether, ABRIM enables researchers to model the advanced imaging parameters and cognitive topologies as a function of age, identify the normal range of values of such parameters, and to further investigate the diverse mechanisms of reserve and resilience.

## Acknowledgments

We express our gratitude to all ABRIM participants for their valuable contributions and dedication of their time. The study was funded by the European FP7 program, FP7-PEOPLE-2013-ITN, Marie-Curie Action, “Initial Training Networks” named “Advanced Brain Imaging with MRI” (no. 608123).

