## Appendix for "The Advanced BRain Imaging on ageing and Memory (ABRIM) data collection: Study protocol and rationale"

*Corresponding author

¶These authors contributed equally.

### Tables

#### S1 Table. Overview variables in ABRIM relative to other databases

| Dataset name | Age range | Participants | Cognitive assessment | Actigraphy | T1-weighted | T2-weighted | MP2RAGE | DWI | T2* or GRE | Resting-state fMRI |
|---|---|---|---|---|---|---|---|---|---|---|
| Cambridge Centre for Ageing and Neuroscience (Cam-CAN) | 18-87 | ~650 | X | | X | X | | X | | X |
| MPI-Leipzig Mind-Brain-Body | 20-35; 59-77 | 227; 74 | X | | | X | X | X | X | X |
| Neurocognitive aging data release | 18-34; 60-89 | 181; 120 | X | | X | | | | | X |
| The Lifespan Human Connectome Project in Aging | 36-100+ | 725 | X | | X | X | | X | | X |
| The National Institute of Health (NIMH) Intramural Healthy Volunteer Dataset | 18-72 | 155 | X | | X | X | | X | X | X |
| UK Biobank | 40-70 | ~50,000 | X | X | X | | | X | X | X |
| Whitehall II Imaging Sub-study | 60-85 | 775 | X | | X | X | | X | | X |
| Lothian Birth Cohort 1936 | ±70 | 866 | X | | X | X | X | X | X | |

| | | | | | | | | | | |
|---|---|---|---|---|---|---|---|---|---|---|
| The Advanced BRain Imaging on ageing and Memory | 18-80 | 295 | X | X | X | X | X | X | X | X |

**S2 Table. Sample characteristics of full sample, per age decade and sex.**

| | | Full sample | 18-30 years | 31-40 years | 41-50 years | 51-60 years | 61-70 years | 71-80 years |
|---|---|---|---|---|---|---|---|---|
| **Full sample** | N | **295** | **46** | **45** | **48** | **51** | **53** | **52** |
| | Age, median (IQR) | **52 (36-66)** | **25 (22-28)** | **35 (33-38)** | **47 (44-49)** | **55 (53-57)** | **65 (62.50-68)** | **73 (72-76)** |
| | Low education, N (%) | **36 (13%)** | **3 (8.3%)** | **4 (9.1%)** | **4 (8.5%)** | **6 (12.8%)** | **9 (17.6%)** | **10 (19.2%)** |
| | Medium education, N (%) | **90 (32.6%)** | **14 (38.9%)** | **18 (40.9%)** | **15 (31.9%)** | **17 (36.2%)** | **16 (31.4%)** | **10 (19.2%)** |
| | High education, N (%) | **151 (54.7%)** | **19 (52.8%)** | **22 (50%)** | **28 (59.6%)** | **24 (51.1%)** | **26 (51%)** | **32 (61.5%)** |
| **Females** | N | **157** | **25** | **25** | **25** | **25** | **27** | **30** |
| | Age, median (IQR) | **52 (36.5-66.5)** | **23 (23-26)** | **36 (33-38)** | **47 (44-49)** | **55 (53-58)** | **65 (62-68)** | **73 (72-76)** |
| | Low education, N (%) | **22 (15.1%)** | **1 (5.3%)** | **3 (12.5%)** | **2 (8.3%)** | **3 (13%)** | **7 (26.9%)** | **6 (20%)** |
| | Medium education, N (%) | **49 (33.6%)** | **8 (42.1%)** | **10 (41.7%)** | **7 (29.2%)** | **10 (43.5%)** | **7 (26.9)** | **7 (23.3%)** |
| | High education, N (%) | **75 (51.4%)** | **10 (52.6%)** | **11 (45.8%)** | **15 (62.5%)** | **10 (43.5%)** | **12 (46.2%)** | **17 (56.7%)** |
| **Males** | N | **138** | **21** | **20** | **23** | **26** | **26** | **22** |
| | Age, median (IQR) | **52 (35.75-66)** | **27 (24-29)** | **33 (32-37.75)** | **46 (43-48)** | **55 (52.75-57)** | **66 (63.50-68)** | **73.50 (72-76)** |
| | Low education, N (%) | **14 (10.7%)** | **2 (11.8%)** | **1 (5%)** | **2 (8.7%)** | **3 (12.5%)** | **2 (8%)** | **4 (18.2%)** |

| | | | | | | | | |
|---|---|---|---|---|---|---|---|---|
| | Medium education, N (%) | **41 (31.3%)** | **6 (35.3%)** | **8 (40%)** | **8 (34.8)** | **7 (29.2%)** | **9 (36%)** | **3 (13.6%)** |
| | High education, N (%) | **76 (58%)** | **9 (52.9%)** | **11 (55%)** | **13 (56.5%)** | **14 (58.3%)** | **14 (56%)** | **15 (68.2%)** |

Data on educational attainment were not available for n = 18 participants (6.4% of the total sample). For females and males, the respective numbers were n = 11 (7% of all females) and n = 7 (5.1% of all males).

**S3 Table. Characteristics of actigraphy sample.**

| | Full sample | 18-30 years | 31-40 years | 41-50 years | 51-60 years | 61-70 years | 71-80 years |
|---|---|---|---|---|---|---|---|
| N | 120 | 13 | 11 | 16 | 26 | 28 | 26 |
| Age, median (IQR) | 57 (44.3-69) | 26 (24-30) | 33 (31-37) | 45.50 (44-48.75) | 54.50 (52-57) | 65 (62-68) | 73 (72-76) |
| Females, N (%) | 68 (56.6%) | 10 (76.9%) | 7 (63.6%) | 8 (50%) | 12 (46.2%) | 17 (60.7%) | 14 (53.8%) |
| Low education, N (%) | 14 (12.2%) | 0 (0%) | 1 (9.1%) | 1 (6.3%) | 1 (4.2%) | 5 (19.2%) | 6 (23.1%) |
| Medium education, N (%) | 30 (26.1%) | 6 (50%) | 3 (27.3%) | 4 (25%) | 9 (37.5%) | 4 (15.4%) | 4 (15.4%) |
| High education, N (%) | 71 (61.7%) | 6 (50%) | 7 (63.6%) | 11 (68.8%) | 14 (58.3%) | 17 (65.4%) | 16 (61.5%) |

Data on educational attainment was not available for n = 5 participants (4.2%).

**S4 Table. Sample characteristics of behavioural and cognitive assessment only participants.**

| | **Full sample** | **18-30 years** | **31-40 years** | **41-50 years** | **51-60 years** | **61-70 years** | **71-80 years** |
|---|---|---|---|---|---|---|---|
| N | 108 | 37 | 7 | 14 | 22 | 20 | 8 |
| Age, median (IQR) | 48 (24-61) | 22 (21-24.5) | 32 (32-35) | 47 (43-48) | 55 (52-57.25) | 64.5 (62-67) | 72 (71.25-74.75) |
| Females, N (%) | 64 (59.3%) | 24 (64.9%) | 5 (71.4%) | 9 (64.3%) | 9 (40.9%) | 11 (55%) | 6 (75%) |
| Low education, N (%) | 5 (4.7%) | 0 (0%) | 1 (14.3%) | 0 (0%) | 0 (0%) | 2 (10%) | 2 (25%) |
| Medium education, N (%) | 32 (31.1%) | 16 (45.7%) | 4 (57.1%) | 5 (38.5%) | 5 (22.7%) | 2 (10%) | 0 (0%) |
| High education, N (%) | 68 (64.2%) | 19 (54.3%) | 2 (28.6%) | 8 (61.5%) | 17 (77.3%) | 16 (80%) | 6 (75%) |

Data on educational attainment was not available for n = 3 participants (2.8%).

### Figures

**S1 Figure. Overview of ABRIM BIDS folder structure.**

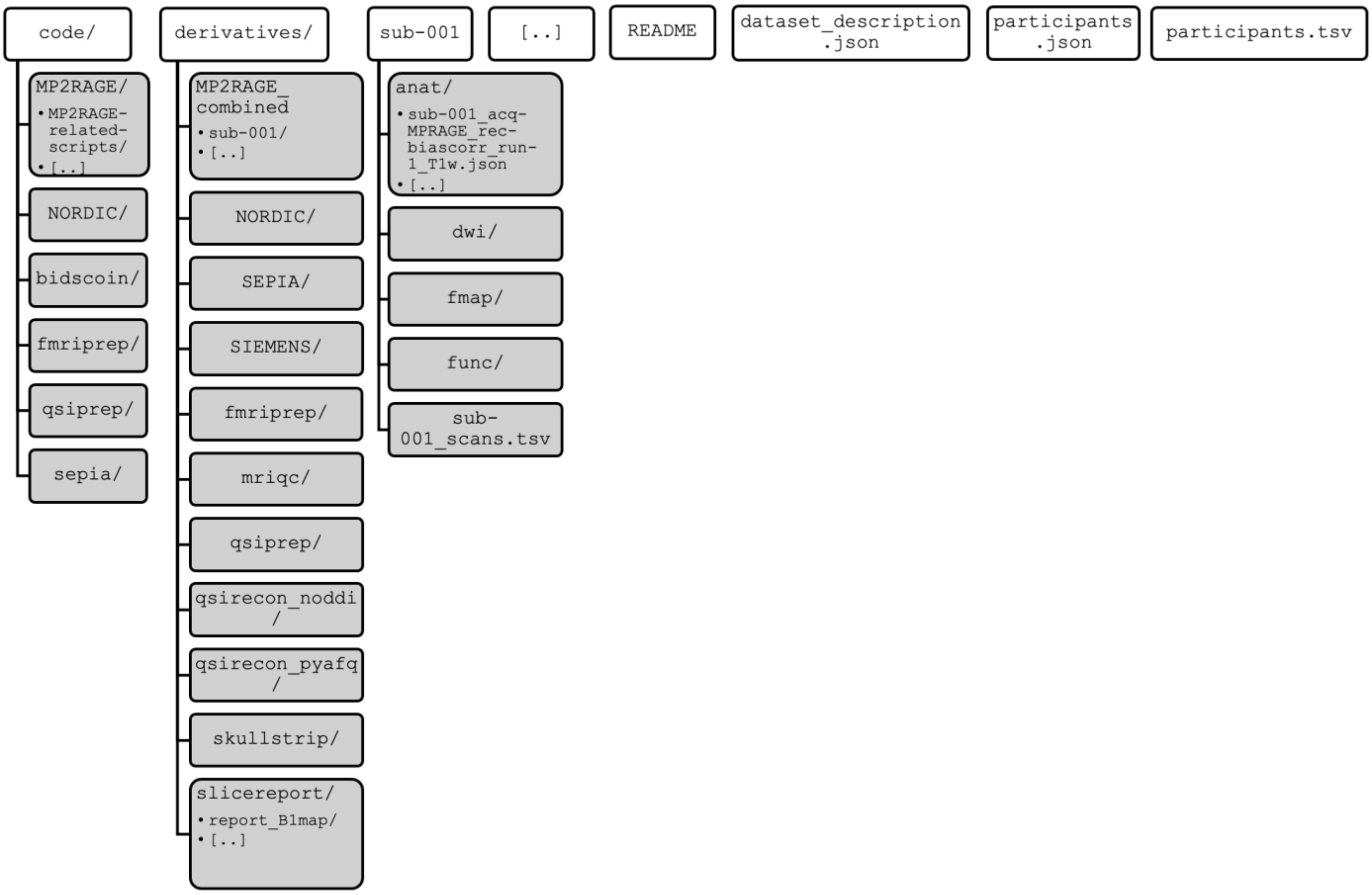

Hierarchical overview of ABRIM MRI data collection folder structure following the Brain Imaging Data Structure (BIDS) standard. The “code” folder contains all scripts that have been applied for data processing. The “derivatives” folder contains all processed and derived data that has been generated from the raw MRI data (e.g., “NORDIC”, “SEPIA”, etc.). Individual subject-specific folders (e.g,. “sub-001”, “sub-002”, etc.) contain modality-specific sub-folders with different types of MRI data (e.g., “anat” contains structural MRI data, whereas “dwi” contains diffusion-weighted imaging data).

**S2 Figure. Group report of T1 images from the MRI Quality Control tool**

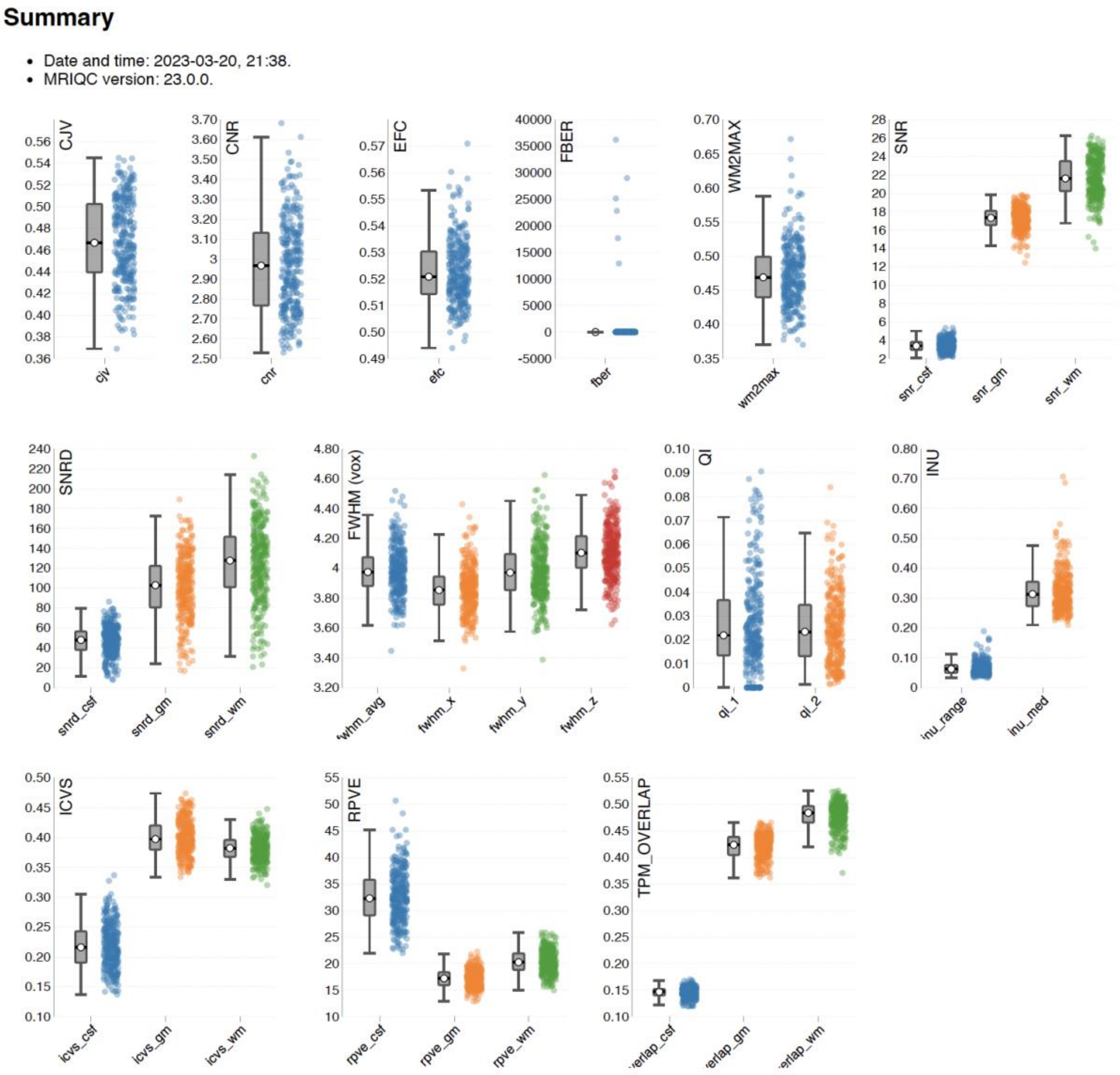

Group anatomical report of T1 scans in ABRIM as generated by the MRI Quality control tool (MRIQC). Contains separate strip-plots for different image quality metrics (IQMs). CJV, coefficient of joint variation; CNR, contrast-to-noise-ratio; EFC, entropy focus criterion; FBER, foreground-to-background energy ratio; WM2MAX white-matter to maximum intenstiy ratio; SNR, signal-to-noise-ratio; SNRD, Dietrich's signal-to-noise-ratio; FWHM (vox), full width half maximum in units of voxels; QI, quality index; INU, intensity non-uniformity; ICVS, intracranial volume fraction; RPVE, residual partial volume effect; TPM_OVERLAP, overlap of issue probability maps of the images and maps from the ICBM nonlinear-asymmetric 2009a template.

**S3 Figure. Group report of T2 images from the MRI Quality Control tool**

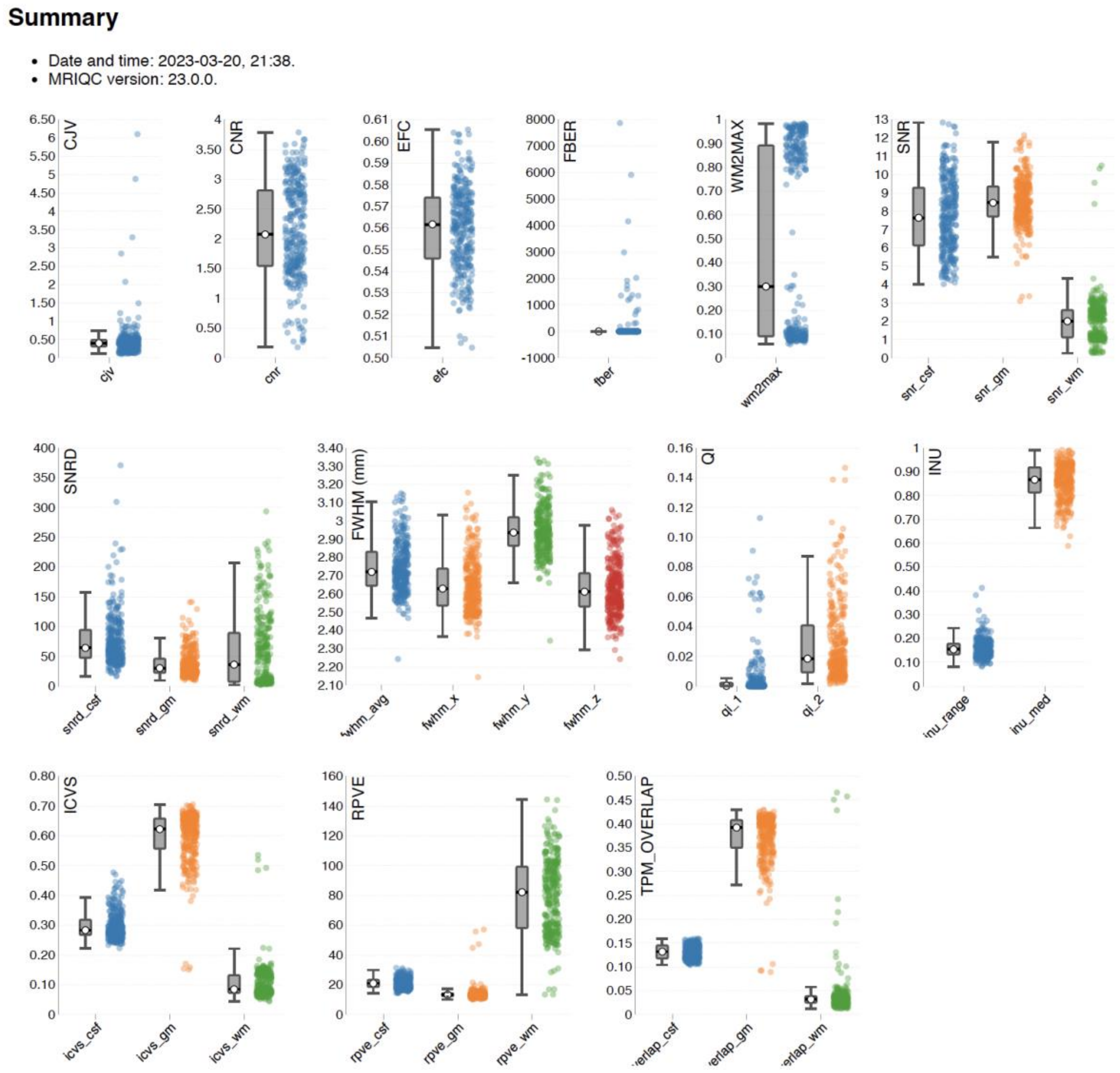

Group anatomical report of T2 scans in ABRIM as generated by the MRI Quality control tool (MRIQC). Contains separate strip-plots for different image quality metrics (IQMs). CJV, coefficient of joint variation; CNR, contrast-to-noise-ratio; EFC, entropy focus criterion; FBER, foreground-to-background energy ratio; WM2MAX, white-matter to maximum intenstiy ratio; SNR, signal-to-noise-ratio; SNRD, Dietrich’s signal-to-noise-ratio; FWHM (vox), full width half maximum in units of voxels; QI, quality index; INU, intensity non-uniformity; ICVS, intracranial volume fraction; RPVE, residual partial volume effect; TPM_OVERLAP, overlap of issue probability maps of the images and maps from the ICBM nonlinear-asymmetric 2009a template.

**S4 Figure. Group report of fMRI images from the MRI Quality Control tool**

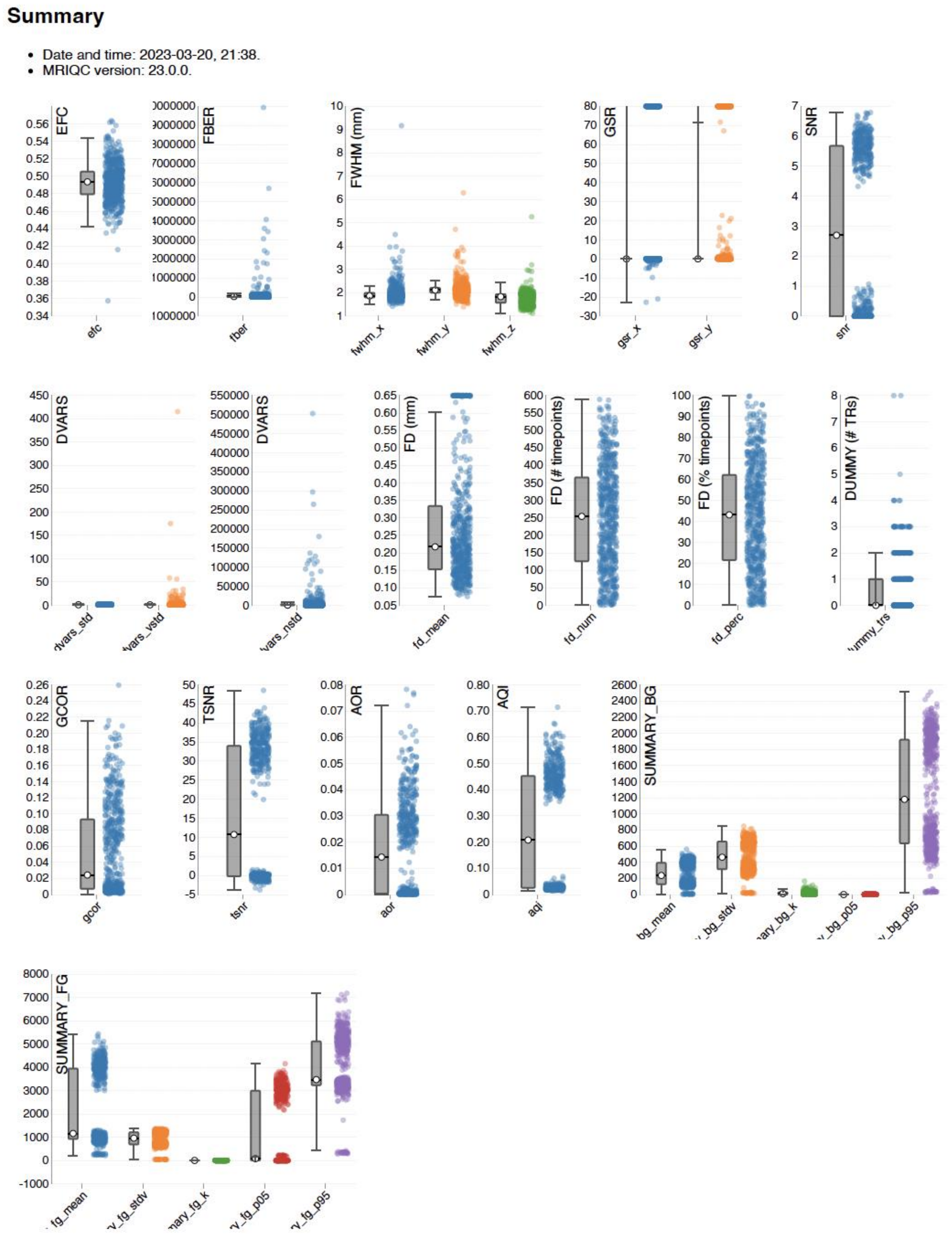

Group anatomical report of T2 scans in ABRIM as generated by the MRI Quality control tool (MRIQC). Contains separate strip-plots for different image quality metrics (IQMs). EFC, entropy focus criterion; FBER, foreground-to-background energy ratio; FWHM, full width half maximum in units of millimetres; GSR, ghost-to-signal ratio; SNR, signal-to-noise ratio; DVARS, index of rate of change of BOLD signal across the entire brain; FD, framewise displacement (number of timepoints and percentage of timepoints above threshold);

DUMMY, number of dummy scans; GCOR, global time-series correlation; TSNR, temporal signal-to-noise ratio; AOR, AFNI's outlier ratio; AQI, AFNI's quality index.

### Appendices

#### S1 Appendix. English translation of memory strategy descriptions.

Many people use techniques or strategies to improve their recall of word combinations. Have you utilized any techniques or strategies to memorize the word pairs you heard? Could you provide a description of these strategies with as much detail as possible below?

*[next page]*

The following list consists of strategies or techniques that people can use to remember word combinations. Read the list of strategies and tick the applicable boxes to indicate which strategies you have employed to enhance your recall of the word pairs. Indicate all strategies that you have used, even if you already described them on the previous page.

| | |
|---|---|
| Concentrate | ☐ |
| Repeat in your head | ☐ |
| Visualize (create imagesin your head) | ☐ |
| Making associations between the words of a pair | ☐ |
| Making a story containing words of a pair | ☐ |
| Making a sentence containing both words of a pair | ☐ |
| Visualize with yourself in a mental image | ☐ |
| Remember specific letters or syllables | ☐ |
| Memorize sounds of word combinations | ☐ |
| Something different, namely………………………………………… | ☐ |
